# 5-Formylcytosine controls nucleosome positioning through covalent histone-DNA interaction

**DOI:** 10.1101/224444

**Authors:** Eun-Ang Raiber, Guillem Portella, Sergio Martínez Cuesta, Robyn Hardisty, Pierre Murat, Zhe Li, Mario Iurlaro, Wendy Dean, Julia Spindel, Dario Beraldi, Mark A. Dawson, Wolf Reik, Shankar Balasubramanian

## Abstract

Nucleosomes are the basic unit of chromatin that ensure genome integrity and control access to the genetic information. The organization of nucleosomes is influenced by the underlying DNA sequence itself, transcription factors or other transcriptional machinery associated proteins and chromatin remodeling complexes (*1–4*). Herein, we show that the naturally occurring DNA modification, 5-formylcytosine (5fC) contributes to the positioning of nucleosomes. We show that the ability of 5fC to position nucleosomes *in vitro* is associated with the formation of covalent interactions between histone residues and 5fC in the form of Schiff bases. We demonstrate that similar interactions can occur in a cellular environment and define their specific genomic loci in mouse embryonic stem cells. Collectively, our findings identify 5fC as a determinant of nucleosomal organization in which 5fC plays a role in establishing distinct regulatory regions that are linked to gene expression Our study provides a previously unknown molecular mechanism, involving the formation of reversible-covalent bonds between chromatin and DNA that supports a molecular linkage between DNA sequence, DNA base modification and chromatin structure.

---

The organization of nucleosomes is an important feature to the chromatin structure. The DNA sequence itself has been shown to be a determinant of nucleosome positioning raising questions about the effect of DNA modifications on chromatin structure. 5-Formylcytosine is generated by the oxidation of 5mC by TET enzymes. It can undergo base excision repair by the thymine DNA glycosylase (TDG) (*5*) and it is therefore thought to mark sites that undergo active demethylation (6). However, we have previously shown that the 5fC distribution was tissue-dependent in mice (7) and that 5fC was a stable modification in genomic DNA *in vivo* (8). Additionally we and others have demonstrated that 5fC can alter the physical properties of the DNA double helix (*9, 10*) and the identification of 5fC-binding chromatin remodelers and transcription factors *in vitro (11, 12*) raises fundamental questions about its involvement in chromatin biology. Moreover, the nature of the formyl group confers specific chemical properties to 5fC that are unique from other cytosine modifications such as 5-hydroxymethylcytosine (5hmC) and 5-carboxycytosine (5caC). We therefore hypothesized that 5fC may be important in nucleosomal organization, particularly during early development, where increased levels of 5fC have been reported (8). To test this hypothesis, we assessed the impact of 5fC on nucleosome core particles formation and stability. We carried out biochemical studies on three different DNA sequences, Widom 601, MMTV-A and 5SrDNA with varying CpG and GC content (Fig. 1A and Supplementary Table 1). All sequences are able to form nucleosomes of different stabilities and properties and have been extensively characterized (13–16). In addition, effects into 5-methylcytosine (5mC) (*17, 18*) and 5-hydroxymethylcytosine (5hmC) (19) on nucleosome occupancy have been investigated on the Widom 601 sequence, however it is unclear how 5fC affects nucleosome formation and stability. To address this, we compared nucleosome occupancy after chaperone-mediated assembly using DNA with unmodified and modified cytosines. We observed that 5mC, 5hmC and 5fC each caused an increase in nucleosome occupancy compared to unmodified Widom 601 DNA (Fig. 1A), with 5fC-DNA significantly (unpaired t-test, p value ≤ 0.0001) displaying the strongest effect (Fig. 1B). Notably we found that only 5fC consistently and significantly (unpaired t-test, p value ≤ 0.01) enhanced nucleosome occupancy on the MMTV-A and 5SrDNA sequences (Fig. 1B and Supplementary Fig. 1A and B).

**Fig. 1.**
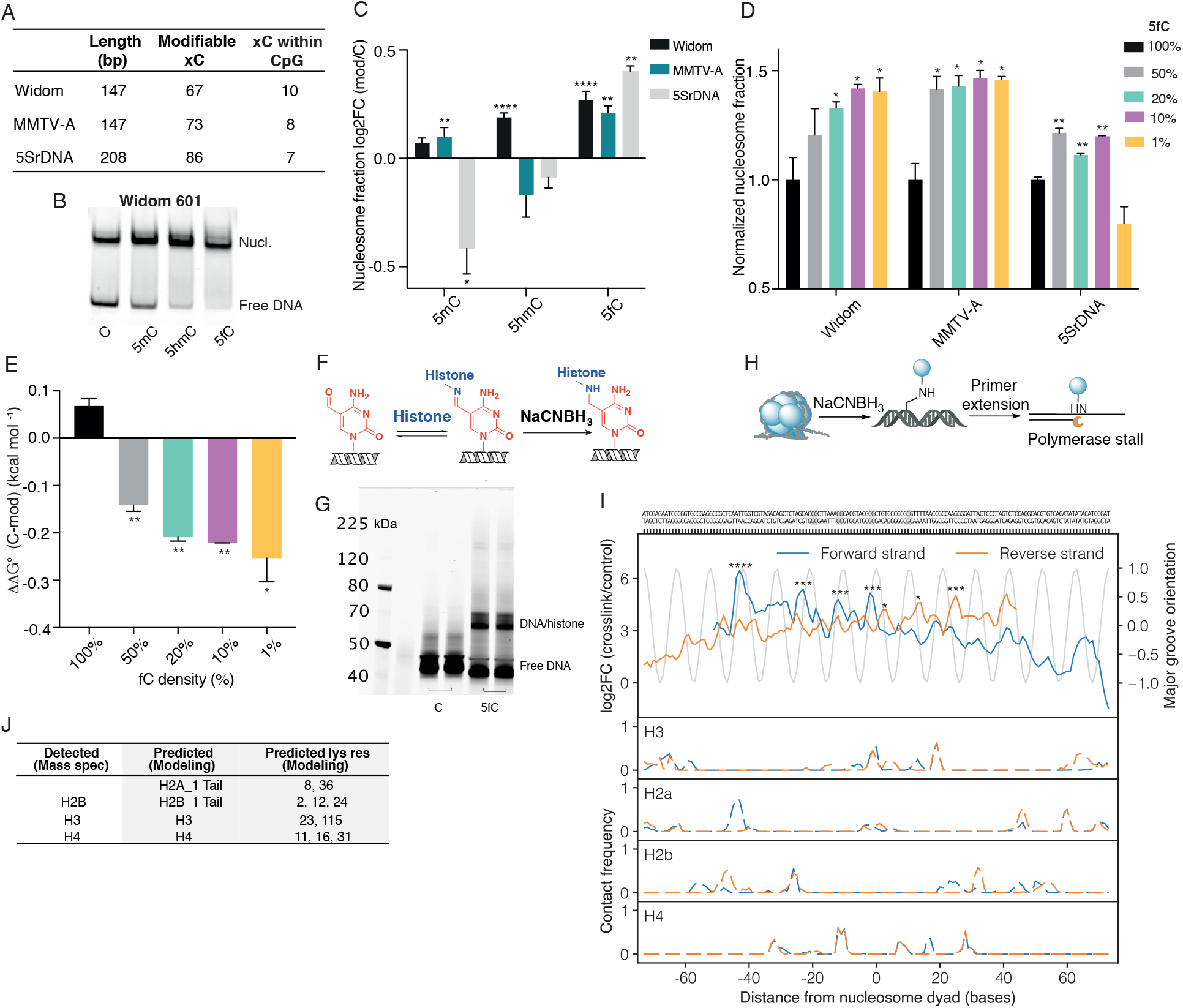
5fC enhances nucleosome occupancy and can form a Schiff base with histone residues. **A**, Double-stranded DNA (Widom 601, MMTV-A and 5SrDNA) comprising unmodified or modified cytosines were used to reconstitute nucleosomes in the presence of NAP1. The table shows the length of the sequences and total number of modifiable cytosines (xC) and number of xC within CpG dinucleotide context. **B**, Non-denaturing polyacrylamide gel shows the nucleosomal (Nucl.) and free DNA fraction of cytosine, 5mC, 5hmC and 5fC Widom DNA. **C**, Log2FC of the nucleosome fraction (Nucl./ total DNA) of cytosine DNA compared to 5mC, 5hmC and 5fC DNA was plotted for the Widom (black), MMTV-A (turquoise) and 5SrDNA sequence (grey). The error bars represent the standard deviation from technical replicates (*n* = 3). Unpaired t test was used to calculate p values (** p ≤ 0.01, **** p ≤ 0.0001). **D**, Nucleosomes were reconstituted using different 5fC density (1–100%) Widom DNA. Normalized nucleosome fractions were plotted against 5fC density. The error bars represent the standard deviation from two technical replicates. **E**, Competitive DNA-histone reconstitution method was used for the measurement of quantitative free-energy of 5fC nucleosomes using the 5S fragment DNA as competitor (4.2 μg and 100 ng Widom). Differences in free energy changes (ΔΔG°) were obtained by subtracting the free energy change for the cytosine Widom DNA from the free energy change for each 5fC density Widom DNA and are represented as the mean along with the standard deviation from two replicates. **F**, 5fC-DNA forms a Schiff base with the primary amines of proximal lysines on histones within the nucleosome, which is then reduced to an irreversible, covalent crosslink using NaCNBH_3_. **G**, 12% Bis-Tris Protein gel electrophoresis was used to separate non-crosslinked from crosslinked nucleosomes. Two replicates are shown for each, cytosine and 5fC (both cy5-labeled). The appearance of a second, higher DNA band was observed for 5fC DNA due to covalent DNA-protein complex that was submitted for mass spec analysis (Supplementary Table 2). **H**, DNA polymerase stalling sites were identified by sequencing to obtain the locations of crosslinked 5fC sites around the histone core. I, Upper panel: The log_2_ fold change between the number of reads stalling in a given position in the polymerase stop assay and the number of reads stalling in a untreated 5fC sample, both for the forward (orange) and reverse strand (blue). The gray sinusoidal line indicates the orientation of the major groove with respect to the histone core, ranging from 1 (histone-core facing) to -1 (facing away from the histone core). Significant (FDR value <0.0001, Benjamini-Hochberg correction on exact p-value from negative binomial distribution) stalling sites (±3 bases) around CpG dinucleotides within the DNA sequence are highlighted in grey. Lower panels: The closest lysines (<5 Å) in histone 2A and B (H2A, H2B), histone 3 (H3) and histone 4 (H4) facing the major grooves of the DNA pointing towards the histone core were identified. Based on the overlap of the computed data with stalling sites (dotted yellow and purple lines for forward and reverse strand) potential 5fC sites involved in the Schiff base formation were identified. J, Table summarizes findings from mass spec analysis (of the higher DNA band from gel), which was supported by data obtained from modeling.

Since the density of 5fC in the fully modified DNA sequences does not reflect densities observed within the genomic DNA context (20) we studied the effect of decreasing 5fC densities (100% -1% 5fC with 1% 5fC ~ 1 5fC per strand) on nucleosome formation. DNA with decreasing 5fC density generally caused increased nucleosome fractions for all three sequences, the only exception being a reduction in nucleosome fraction at 1% 5fC for the 5SrDNA sequence (Fig. 1C). This data was supported by our results obtained from reconstitution by competition dialysis (13) to determine differences in ΔG. Here our results showed that 1% 5fC DNA caused the most favorable shift in free energy change (ΔΔG° = -0.253± 0.05 kcal mol^-1^) for nucleosome formation when referenced to unmodified DNA (Fig. 1D and Supplementary Fig. 1C). The capacity for DNA bending as required by nucleosome structure, is an important determinant to nucleosomal organization and it has been reported that a single 5fC unit significantly increases the flexibility of DNA (10), which may explain our observation.

The aldehyde functionality of 5fC is prone to nucleophilic addition (e.g. by the *ε* - amino group of lysine) suggesting a chemical mechanism for 5fC to promote nucleosome occupancy. It has recently been suggested that 5fC can form a covalent bond with proteins (21, 22). The formation of reversible, covalent Schiff base (imine) bonding between the formyl group of 5fC with proximal lysine residues of histones would provide a selective, covalent mechanism for positioning nucleosomes even with a single 5fC per nucleosome (Fig. 1E). To explore this hypothesis, we employed the reducing agent NaCNBH_3_ on nucleosome core particles obtained from 5fC-containing Widom sequence to irreversibly trap any Schiff base after its formation, (Fig. 1F and Supplementary Fig. 1D). Denaturing gel electrophoresis showed the appearance of a new higher molecular weight band that was not observed in the cytosine containing control DNA (Fig. 1F). Also in the absence of NaCNBH_3_ or treatment of nucleosomes with EtONH_2_ prior reduction (as a competitor to lysine residues) we did not observe a higher molecular weight band, consistent with reversible Schiff base formation between 5fC-DNA and histone proteins (Supplementary Fig. 1E). Proteomics mass spectrometry analysis of the higher molecular weight band from the gel identified the presence of histone core proteins H2B, H3 and H4 in the complex, consistent with covalent DNA adduct formation with these histones (Supplementary Table 2). We then exploited polymerase-stalling, caused by the stable crosslink formed after NaCNBH_3_ reduction, using a single primer extension reaction followed by sequencing to identify specific 5fC-lysine interactions (Fig. 1G and Supplementary Fig. 1F). Particularly, we observed significant stalling sites where 5fC was in a CpG context and in the major groove of DNA facing the histone core demonstrating that formylation of CpG sites, rather than C residues, enhanced covalent interactions between histone proteins to the DNA sequence (Fig. 1H and Supplementary Fig. 1G). (3). It is also noteworthy that significant stalling sites were enriched in the vicinity of the nucleosome dyad. To identify the most probable interactions between histone residues and 5fC, we associated 5fC-induced polymerase stalling sites with the most proximal (within 5 Å) lysine-nucleotide distances computed from molecular dynamics simulations that model the Widom nucleosome (<Fig. 1I and Supplementary Fig. 1G). We observed that lysine residues in close proximity to significant stalling sites in the dyad region were mostly found in the H3 protein (Fig. 1X and Table 1). These data suggest a structural model whereby specific interactions between 5fC and lysine residues from the H3 protein may contribute to nucleosome positioning.

Since the intrinsic sequence preference of nucleosomes itself is a known determinant for the nucleosomal organization (*3, 23*), we next asked whether 5fC-associated preference for nucleosomes had an impact on DNA sequence-directed nucleosome positioning. We generated four indexed libraries of unmodified or fully modified DNA sequences by sonication of the genomic DNA, adapter ligation and subsequent PCR amplification using either dCtp, d5mCtp, d5hmCtp or d5fCtp (Fig. 2A, see Method). These libraries of oligonucleotides were then pooled (1:1:1:1) and subsequently used in a nucleosome reconstitution assay where the total amount of DNA was in excess compared to histone proteins to select for higher affinity DNA sequences. Nucleosomes were separated from free DNA on a non-denaturing gel, followed by extraction of the nucleosome fraction. Sequencing of the input DNA pool (before nucleosome reconstitution) and the nucleosome fraction allowed the identification of significantly (p-value ≤ 0.00001) enriched nucleosomal DNA sequences from each library. We identified 2,977 sequences for cytosine, 884 for 5mC, 10,537 for 5hmC and 2,481 for 5fC DNA libraries. Fig. 2B shows the nucleosome enrichment (nucleosome reads/ reads of the corresponding sequence in the input) for the different DNA libraries revealing that overall 5fC was significantly (two-sided Mann-Whitney test, p value ≤ 0.0001) associated with the highest enrichment compared to unmodified and other cytosine modifications. Our study also revealed that positions of 5fC-associated nucleosomes overlapped the least with nucleosomes associated with cytosine, 5mC or 5hmC demonstrating the ability for 5fC DNA to uniquely position nucleosomes (Fig. 2C). When we calculated the actual fraction of overlapping bases (Fig. 2D) between all libraries (vertical n horizontal), we observed that over 80% of the 5mC-associated nucleosome positions overlapped with cytosine and 5hmC nucleosomes, whereas 5fC (and 5hmC) only shared 20-30% of nucleosome positions between modifications. This observation is depicted in Fig. 2E, where MNase signals were enriched either in the 5fC or cytosine/5mC nucleosomes at two representative genomic loci. We next analysed the GC content and CpG density associated with the DNA sequences enriched in nucleosomes. We found that cytosine and 5mC-associated nucleosome enrichment was largely independent of GC content (Fig. 2F). In contrast, 5fC-associated enrichment was influenced by low GC content. Similarly, lower CpG density was associated with higher enrichment of 5fC-associated nucleosomes (Fig. G) demonstrating some interplay between DNA sequence context and DNA modifications on nucleosome formation in vitro. Taken together, these data show that formylation of DNA sequences with low GC and CpG density uniquely enhances nucleosome affinity.

**Fig. 2.**
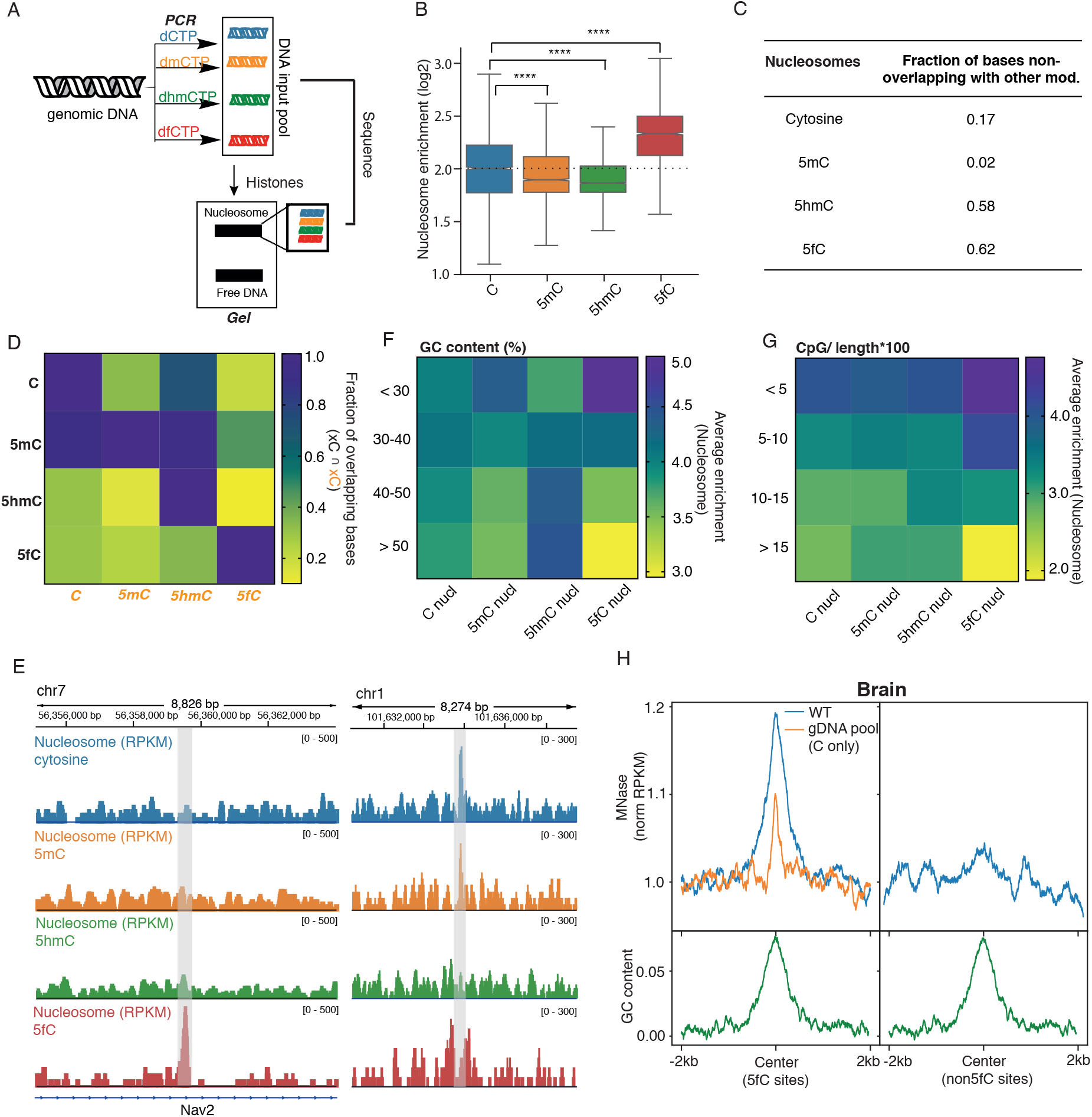
5fC within genomic sequence context enhances nucleosome positioning in vitro. **A**, A pool of DNA sequences containing either cytosine, 5mC, 5hmC or 5fC was generated by PCR using genomic DNA and used for subsequent nucleosome reconstitution and sequencing. **B**, Nucleosome enrichment (nucleosome library/input DNA library) for cytosine (blue), 5mC (orange), 5hmC (green) and 5fC (red) DNA was plotted. Notched boxplot shows the 1st, 2nd (median) and 3rd quartile, with whiskers extending to the minimum and maximum. P-values were obtained using the two-sided Mann-Whitney test (**** p-value ≤ 0.0001). **C**, Table shows the number of unique fraction of bases associated with cytosine, 5mC, 5hmC or 5fC nucleosomes. **D**, Heatmap showing the fraction of overlapping bases when intersecting cytosine, 5mC, 5hmC and 5fC-associated nucleosome sequences with each other. **E**, Representative genomic loci showing unique 5fC-associated nucleosome positioning (left panel, red) and overlap between cytosine (blue) and 5mC-associated (orange) nucleosome positioning (right panel). **F, G**, Heatmap showing sequence features linked to nucleosome enrichment. **H**, MNase reads (normalized RPKM) of reconstituted nucleosomes using genomic DNA extracted from mouse embryonic hindbrain (11.5 days) show enrichment around 5fC brain sites (left panel, blue line (WT)). MNase signals around 1) non-5fC sites (right panel, blue line) with comparable GC content (green line) and around 2) 5fC sites when genomic DNA containing only cytosine was used for reconstitution (WT PCR C only, left panel, orange line) show depletion of MNase signals.

We next investigated the effects of 5fC within its natural sequence context, i. e. mainly within CpGs context, and density on nucleosome formation. We therefore used endogenous mouse embryonic brain genomic DNA to assemble nucleosomes and subsequently performed MNase-sequencing. Fig. 2H (left panel) represents normalized MNase reads at 5fC sites previously identified in hindbrain (7) and 2kb up- and downstream of these sites. Our data revealed that natural 5fC sites are characterized by increased nucleosome density, as judged nucleosome enrichment, consistent with our biochemical observations using synthetic DNA model templates. In contrast, non-5fC sites with similar GC content did not show any enrichment in nucleosome density (Fig. 1H, right panel). To demonstrate that the observed impact of 5fC on nucleosome organization was independent of DNA sequence preference of nucleosomes, we showed that removal of 5fC by generating genomic DNA only containing cytosines (by “erasing” 5fC through PCR amplification) resulted in the depletion of nucleosomes at the same 5fC sites (Fig. 1H, left panel). Collectively, our data show the ability of 5fC to position nucleosomes within the genomic DNA sequence context, both at unnatural (and fully modified) and at naturally (low 5fC density) occurring 5fC genomic loci.

Since 5fC distribution is tissue-specific (7) and its level is dynamic throughout development (8) it raises the question if the ability of 5fC to position nucleosomes is involved in the tissue-specific nucleosomal organization. We therefore generated genome-wide nucleosome maps in hindbrain and heart (E11.5) using MNase-seq to investigate the impact of 5fC on nucleosome organization *in vivo.* We observed clear nucleosome positioning centred at 5fC sites (7) in both, hindbrain and heart tissues (Fig. 3A), consistent with our *in vitro* observations. Our study also showed that on average nucleosome occupancy at 5fC sites was significantly higher (p-value ≤ 0.0001, two-sided Mann-Whitney U-test) than at all other nucleosome positions, both in heart and hindbrain, demonstrating an intrinsic 5fC DNA preference of nucleosomes that support a role for 5fC in determining the organization of nucleosomes *in vivo* (Supplementary Fig. 2A). When we looked into the sequence context of naturally occurring 5fC sites in brain, we found that increasing 5fC levels were associated with increasing CpG density (Supplementary Fig. 2C). Nucleosomes are generally found to be depleted at CpG dense regions such as CpG islands (CGIs) *in vivo,* with a clear link between CpG content and nucleosome depletion (24, 25). Notably, we observed that 5fC-containing CGIs were actually enriched in nucleosomes compared to CGIs that lack 5fC. We also found that in contrast to non-5fC CGIs, mainly found at promoters, the subset of 5fC-CGIs associated with nucleosomes was located in gene bodies of actively transcribed genes fundamental for brain development including Zfp238 (26), Tcf3 (27), Mycn (28), Foxp4 (29), Spen (*30*) and Mid1 (31), highlighting the importance of 5fC in the establishment of distinct nucleosomal organization at tissue-specific active genes. Given our previous observation that 5fC was enriched (7) at sites marked by H3K27ac and H3K4me1, both hallmarks of active enhancers, we next investigated the effect of 5fC on chromatin structure at these sites. For both, H3K27ac and H3K4me1, we observed higher nucleosome density in the presence of 5fC compared to non 5fC sites revealing a differential nucleosomal landscape at regulatory loci containing 5fC (Fig. 3F and G). Enhancers and promoters marked by histone modifications including H3K4me1 and H3K27ac are generally nucleosome depleted (4). In contrast, some active tissue-specific enhancers, characterized by relatively high nucleosome occupancy and accessibility that bind pioneer transcription factors to nucleosomal DNA, are believed to stimulate transcription (*32, 33*). Taken together our data support a model where 5fC is involved in the organization of nucleosomes at regulatory regions important for brain-specific activity. This hypothesis would suggest that 5fC-associated nucleosome positioning could be linked to enhanced, brain-specific gene expression. To evaluate this, we compared the expression of all genes linked to predicted enhancers (*34*) with genes linked to predicted enhancers containing 5fC-associated nucleosomes (Fig. 3H). We found that genes linked to 5fC-nucleosome enhancers were indeed significantly more highly expressed (p-value ≤ 0.0001, two-sided Mann-Whitney U-test) supporting a role of 5fC in the organization of nucleosomes at regulatory sites that is important for gene regulation.

**Fig. 3.**
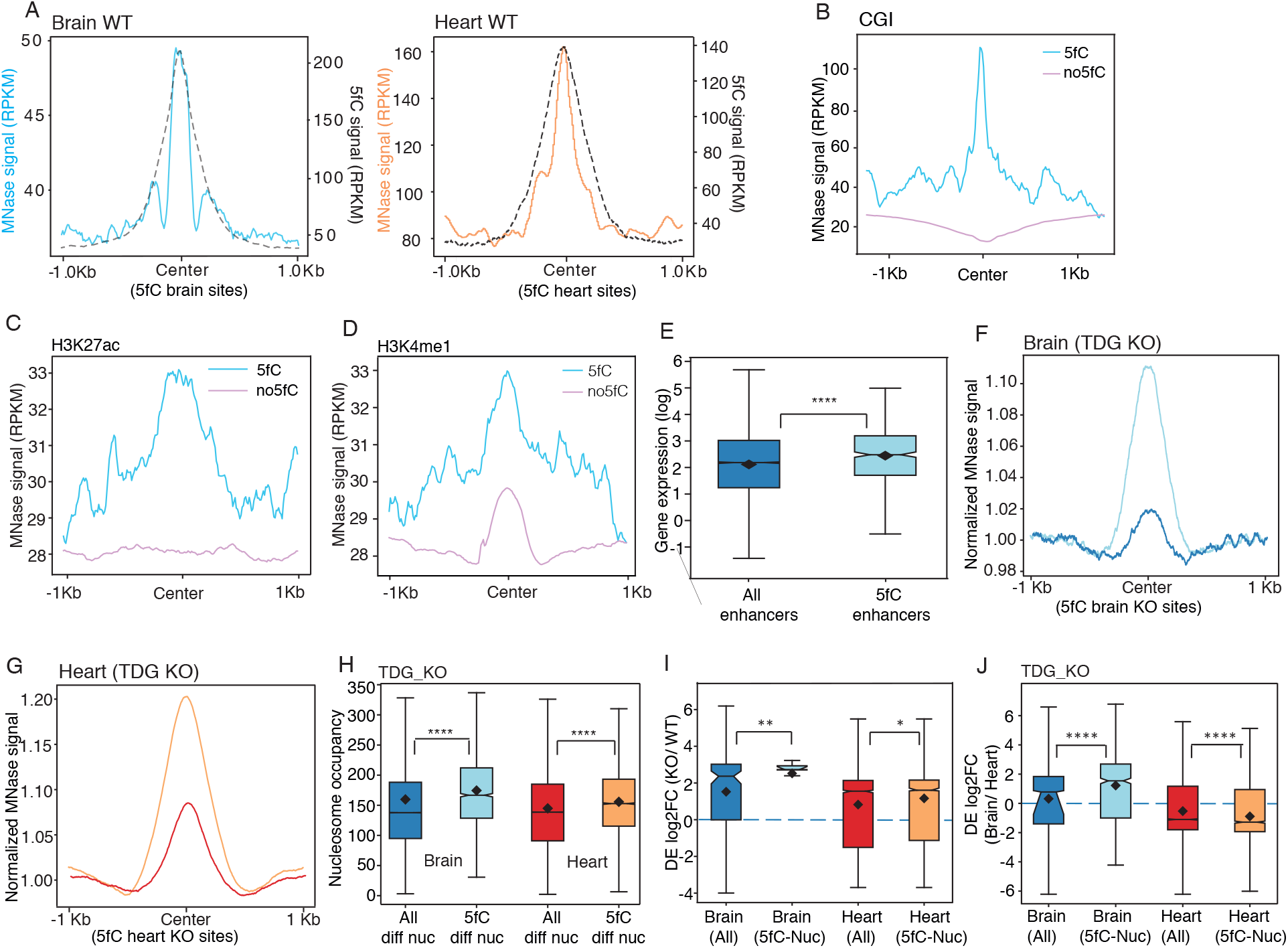
*5fC is a determinant of nucleosomal organization in vivo* that is linked to gene expression. **A**, Normalized MNase signal was plotted around 5fC sites (dotted black line) in hindbrain (blue) and heart (orange) revealing positioning of nucleosomes at the center of 5fC sites. **B**, MNase signal (RPKM) around 5fC CGI (blue lines) were compared to non-5fC CGI (orange) in hindbrain showing that 5fC at CGIs can dictate nucleosome enrichment. **C, D**, H3K27ac and H3K4me1 sites also show differential nucleosome density that is dependent on 5fC. The figure illustrates the enrichment of nucleosomes at H3K27ac and H3Kme1 sites that overlap with 5fC sites. **E**, Comparison of gene expression (log RPKM) of a subset of enhancers marked by 5fC-nucleosomes (*n*= 517, light blue) and all enhancer sites (*n*= 13286, dark blue) in WT hindbrain. Notched boxplot show that the presence of 5fC at enhancer sites is correlated with significantly higher gene expression (p-value ≤ 2.e-8, two-sided Mann-Whitney U-test) of their associated genes compared to all expressed genes (predicted enhancer-gene list from Shen et al, Nature 2007(35)). Boxplot shows 1st, 2nd (median) and 3rd quartile, with notches representing the confidence interval around the median and the black diamond the mean, and the whiskers indicate the reach of the data points beyond the 1st (Q1) and 3rd (Q3) quartile (e.g. Q1+1.5*(Q3-Q1)). **F, G**, Normalized MNase signal around 5fC KO and non-5fC sites with comparable GC content in hindbrain (light blue for 5fC and dark blue for non-5fC) and heart tissues (orange for 5fC and red for non-5fC) show that 5fC sites are enriched in nucleosomes also in KO tissues. H, Notched boxplot shows nucleosome occupancy of differentially positioned nucleosomes in TDG KO hindbrain. Differentially positioned nucleosomes (TDG KO / WT) were identified by DANPOS (FDR < 0.05, Benjamini-Hochberg correction on Poisson p-value). Average nucleosomes occupancy of 5fC-associated differentially positioned nucleosomes (*n*= 2196 for brain and n= 9967 for heart) was significantly (p-value < 0.0001, two-sided Mann-Whitney U-test) higher than all differentially positioned nucleosomes (*n*= 357079 for brain and n= 439541 for heart). **I**, Notched boxplot show that 5fC-associated nucleosomal reorganization at enhancer sites in TDG KO (*n*= 40 for brain and n= 220 for heart) is linked with significantly increased gene expression in hindbrain (blue, p-value < 0.001, two-sided Mann-Whitney U-test) and heart (orange, p-value < 0.05, two-sided Mann-Whitney U-test) tissue compared to all differentially expressed genes (*n*= 87 for brain and n= 188 for heart). **J**, Differential expression analysis between hindbrain (blue) and heart (orange) tissues in TDG KO. Significantly higher (p-value < 0.0001, two-sided Mann-Whitney U-test) differential expression was observed between hindbrain and heart tissue of genes linked to enhancers containing 5fC-associated nucleosomes (for brain *n*= 422 and for heart *n*= 761) than all enhancers (for brain n= 1827 and for heart n= 2082). All boxplots in this figure show the 1st, 2nd (median) and 3rd quartiles, with notches representing the confidence interval around the median and the black diamond the mean, and the whiskers indicate the reach of the data points beyond the 1st (Q1) and 3rd (Q3) quartile (e.g. Q1+1.5*(Q3-Q1)).

To further test this hypothesis, we used the TDG KO model to study the consequences of 5fC redistribution. The DNA repair enzyme TDG can excise 5fC and 5caC (5), and consequently genetic deletion of TDG results in the increase of 5fC levels with up to 10-fold enhancement of 5fC sites genome-wide (7) concomitant with “new” 5fC sites in embryonic hindbrain and heart tissues at embryonic day 11.5 (E11.5). When we examined the consequences of TDG-dependent enrichment of 5fC sites on the nucleosomal organization, we observed positioning of nucleosomes centered at 5fC sites in hindbrain and heart, similar to wild type, whereas nucleosomes were depleted at sites that lacked 5fC sites (Fig. 3I and 3J). When we analysed the differential nucleosome occupancy between WT and TDG-KO we found significantly higher nucleosome occupancy (p-value ≤ 0.0001, two-sided Mann-Whitney U-test) at sites where 5fC had been acquired, both in hindbrain and heart (Fig. 3K). To investigate a potential link between differential nucleosomal organization at emergent 5fC sites in the TDG KO and changes in gene expression, we considered differential expression of genes for which there was information on their predicted enhancers (Fig. 3L). Increased gene expression (p-value ≤ 0.01, two-sided Mann-Whitney U-test) was observed for genes where differential 5fC-associated nucleosomes aligned at the enhancers. Finally, we compared tissue-specific gene expression in WT and TDG KO tissues. We observed that particularly in WT hindbrain, differential gene expression (hindbrain versus heart) was significantly higher (p-value < 0.05, two-sided Mann-Whitney U-test) for genes linked to differential 5fC-nucleosomes enhancers (present in hindbrain but not heart) (Supplementary Fig. 2D). Notably, in the TDG KO we found that many more differential 5fC-nucleosomes now overlapped with enhancers, 761 in heart and 422 hindbrain, that were also linked to significant (p-value < 0.0001, two-sided Mann-Whitney U-test) differential gene expression of their associated genes (Fig. 3M) demonstrating the importance of 5fC-associated nucleosome organization.

Now that we have shown that 5fC can interact with histone residues *via* the formation of Schiff base and that 5fC-associated nucleosome positioning at regulatory sites *in vivo* was important, we next evaluated whether this covalent 5fC and histone interactions may provide a molecular mechanism in native chromatin context. We therefore chemically trapped (using NaCNBH_3_) any Schiff base formed in nuclei from TDG KO mESC that contain relatively high levels of 5fC (35). Subsequent covalent histone-DNA complexes were isolated by histone chromatin immunoprecipitation (ChIP) using four different antibodies to H1, H2A, H3 and actin (Fig. 4A). Non-covalent DNA/ protein interactions were disrupted by guanidine HCl treatment followed by extensive washing through a filter column. We assessed the enrichment of DNA libraries obtained from ChIP using the reduced (NaCNBH_3_-treated) over untreated sample by qPCR. We observed significant enrichment (t-test, p value ≤ 0.01) when the antibody for histone 3 (H3) was used, but not H1 or H2A, suggesting covalent Schiff base formation between 5fC and H 3 (Fig. 4B). We subsequently sequenced two biological replicates of the H3 ChIP libraries to characterize the sites of Schiff base formation. We found a total of 2,693 peaks (across two replicates) that were correlated with existing 5fC maps (*36*) to identify the distribution of covalently 5fC-bound histones in the genome. We found 370 sites that overlapped with 5fC sites, half of which were found within genes (164 genes). Figure 4C shows a representative genomic locus, where 5fC and H3 sites overlaped within the gene body. To further determine functional consequences of H3/5fC interactions within genes, we examined engaged Pol II sites using global nuclear run-on coupled with deep sequencing (GRO-seq) datasets (using datasets from Wang et al. (*37*)) of mESCs TDG KO. Metagene analysis of the Gro-Seq signals in mESCs TDG KO after 0, 10, 20 and 30 minutes of synchronized transcription showed slowing down of Pol II transcription elongation at genes where we have mapped H3-5fC interaction sites (“crosslinked”) compared to “all”, 5fC (without crosslink) and 5caC containing genes with a new wave of transcription starting before 80 kb (Fig. 4 D). Notably, we found that crosslinked H3 sites were also enriched before 80 kb. Finally, our analysis revealed an accumulation of transcriptionally engaged Pol II within 2 kb downstream from the center of all crosslinked H3 sites (Fig. 4E) suggesting involvement of covalent 5fC-H3 interactions in transcription regulation.

**Fig. 4.**
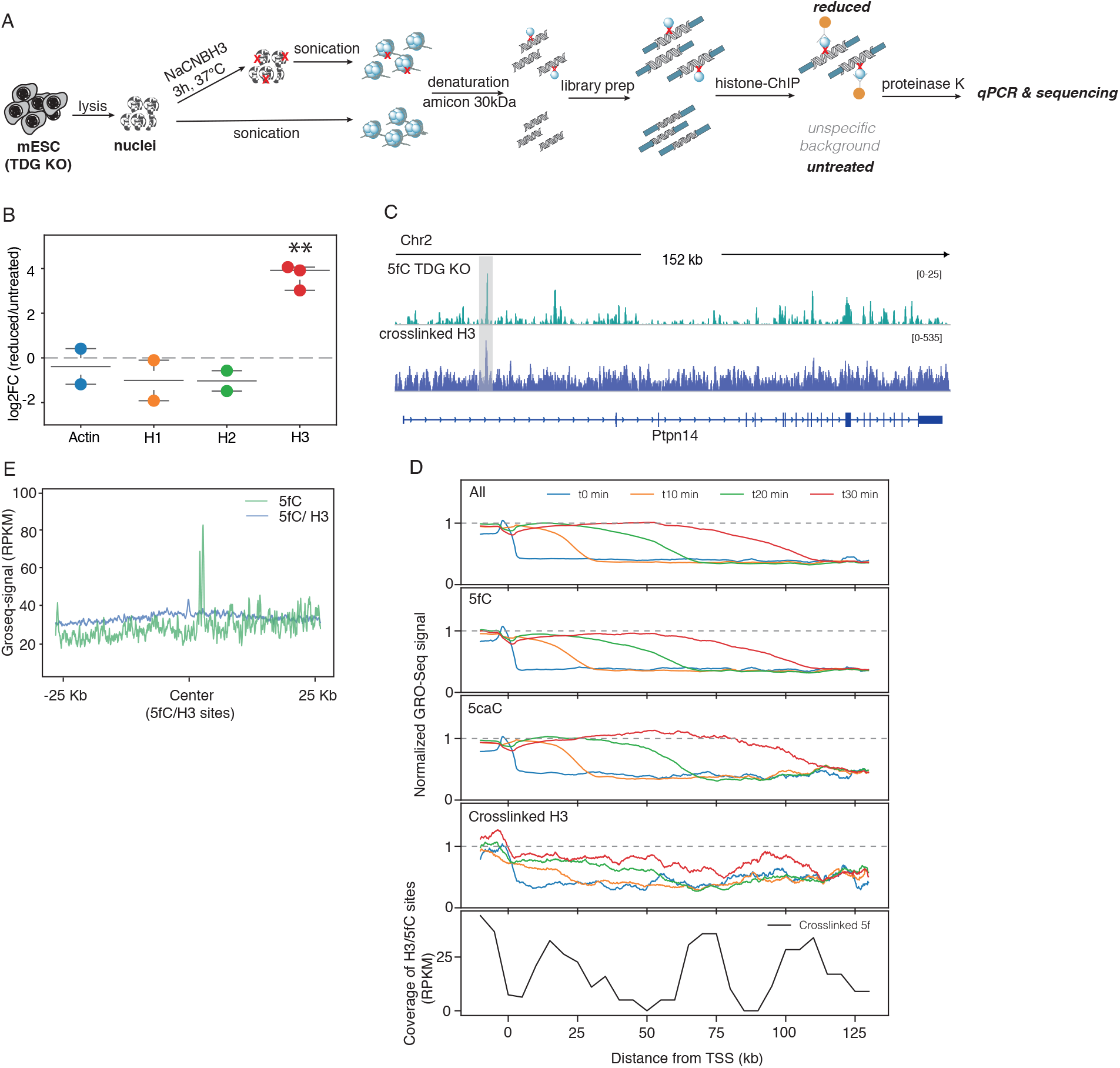
5fC can form Schiff base with histones in chromatin context that impacts polymerase activity. **A**, Workflow for the detection of *in vivo* Schiff base in mESC TDG KO. Key steps involve the reduction of the imine bond using NaCNBH_3_ followed by denaturation of proteins to disrupt any non-crosslinked DNA-protein interaction and subsequent histone ChIP, with no reduction for the control. **B**, Log2FC of reduced/untreated samples after ChIP-qPCR reveal significant enrichment (t-test, ** p-value ≤ 0.01) of H3 immunoprecipitated chromatin after reduction. Scatter dot plot show the values for individual replicates for actin (blue), H1 (orange), H2A (green) and H3 (red) with lines indicating the mean with standard deviation. **C**, Representative genomic loci showing the overlap between 5fC and 5fC/H3 sites in mESC TDG KO. **D**, Metagene analysis of normalized Gro-Seq signal at genes containing All, 5fC, 5caC or crosslinked H3 sites at 0 (blue), 10 (orange), 20 (green) and 30 minutes (red) after DRB treatment. Bottom panel shows the distribution of crosslinked H3 coverage (RPKM) across the genes **E**, Gro-Seq signals (RPKM) centered around 25kb up- and downstream of H3/5fC sites (green) and 5fC only (blue) sites.

In summary, we have shown that 5fC is a determinant of nucleosome organization *in vitro* and *in vivo.* Changes in 5fC patterns caused by loss of TDG (KO) led to nucleosome re-positioning and occupancy linked to changes in gene expression. We demonstrated *in vitro* that 5fC can form a covalent, reversible Schiff base within the nucleosome context that may explain the intrinsic preference of nucleosomes for 5fC DNA. Our data in mESCs provide evidence that 5fC and histone residues may also interact covalently in a cellular context. These 5fC-H3 sites may provide a stalling site for Pol II for the recruitment of proteins involved in transcription elongation or chromatin remodeling. Collectively our data support a model whereby 5fC contributes to the positioning of cell and tissue-specific nucleosomes providing a molecular mechanism to help explain how 5fC regulates gene expression during development and in the reinforcement of cell identity.

## Acknowledgments

The Balasubramanian laboratory is supported by core funding from Cancer Research UK (C14303/A17197). SB is a Senior Investigator of the Wellcome Trust (grant no. 099232/z/12/z). ZL is supported by A*STAR (Singapore). The Reik laboratory is supported by BBSRC (BB/ K010867/1) and the Wellcome Trust (095645/Z/11/Z).

